# The genome-wide transcriptional response to varying RpoS levels in *Escherichia coli* K-12

**DOI:** 10.1101/082537

**Authors:** Garrett T. Wong, Richard P. Bonocora, Alicia N. Schep, Suzannah M. Beeler, Anna J. Lee Fong, Lauren M. Shull, Lakshmi E. Batachari, Moira Dillon, Ciaran Evans, Carla J. Becker, Eliot C. Bush, Johanna Hardin, Joseph T. Wade, Daniel M. Stoebel

## Abstract

The alternative sigma factor RpoS is a central regulator of a many stress responses in *Escherichia coli.* The level of functional RpoS differs depending on the stress. The effect of these differing concentrations of RpoS on global transcriptional responses remains unclear. We investigated the effect of RpoS concentration on the transcriptome during stationary phase in rich media. We show that 23% of genes in the *E. coli* genome are regulated by RpoS level, and we identify many RpoS-transcribed genes and promoters. We observe three distinct classes of response to RpoS by genes in the regulon: genes whose expression changes linearly with increasing RpoS level, genes whose expression changes dramatically with the production of only a little RpoS (“sensitive” genes), and genes whose expression changes very little with the production of a little RpoS (“insensitive”). We show that sequences outside the core promoter region determine whether a RpoS-regulated gene in sensitive or insensitive. Moreover, we show that sensitive and insensitive genes are enriched for specific functional classes, and that the sensitivity of a gene to RpoS corresponds to the timing of induction as cells enter stationary phase. Thus, promoter sensitivity to RpoS is a mechanism to coordinate specific cellular processes with growth phase, and may also contribute to the diversity of stress responses directed by RpoS.

**Importance:** The sigma factor RpoS is a global regulator that controls the response to many stresses in *Escherichia coli.* Different stresses result in different levels of RpoS production, but the consequences of this variation are unknown. We describe how changing the level of RpoS does not influence all RpoS-regulated genes equally. The cause of this variation is likely the action of transcription factors that bind the promoters of the genes. We show that the sensitivity of a gene to RpoS levels explains the timing of expression as cells enter stationary phase, and that genes with different RpoS sensitivities are enriched for specific functional groups. Thus, promoter sensitivity to RpoS is a mechanism to coordinate specific cellular processes in response to stresses.

## Introduction

Genome-wide measurements of RNA levels have revolutionized our understanding of how cells organize their patterns of transcription. These studies have given us snapshots of how patterns of gene expression change in response to changes in the external environment. They have also allowed us to define the regulons controlled by specific transcription factors. A major weakness of the vast majority of these studies is that they explore the function of a regulatory protein only by comparing expression of target genes in a wild-type strain to a gene knock-out, or to a mutant with a single diminished or increased level of activity. While some genetic regulatory networks are certainly switch-like (1, 2) and can be fully characterized by only two levels of activity, many other regulatory proteins vary continuously in their abundance and/or activity. How regulons respond to a range of regulator levels is a largely unstudied question (3). Should we expect all genes in a regulon to increase or decrease expression by the same relative amount following a change in abundance of a regulatory protein? Or should we expect genes to respond in different ways? These questions motivate this study.

A paradigmatic example of a bacterial regulatory protein whose abundance and activity vary continuously in response to different conditions is the alternative sigma factor RpoS in *Escherichia coli.* Transcription by RNA polymerase containing RpoS is responsible for the general stress response (4–6). Under conditions of optimal growth in the laboratory (such as exponential phase growth in rich media at 37°C), RpoS levels are nearly undetectable in the model *E. coli* strain K-12. As conditions become poorer for growth, either because cells begin to starve for various nutrients, or because they face physical challenges such as low temperature or elevated osmolarity, RpoS levels rise (4–6). RpoS coordinates the transcription of genes that are critical for the response to these stresses.

RpoS expression is not an all-or-none phenomenon. For example, RpoS levels rise continuously as cells transition from exponential growth to stationary phase (7). Moreover, starvation for different nutrients upregulates RpoS to differing levels (8, 9). This level of control over RpoS levels is accomplished by regulating transcription, translation, and protein degradation (4, 7, 10), allowing for careful control over protein levels. In addition to regulation of protein abundance, RpoS activity can also be directly modulated by a number of factors, such as Crl (11).

Not only do RpoS levels vary across conditions for a single strain, but different strains of *E. coli* also differ in their patterns of expression of RpoS. For example, naturally occurring strains can differ in the amount of RpoS they produce during exponential phase (12) or stationary phase (13). All studies that have measured RpoS levels in naturally occurring strains of *E. coli* have detected variation (13–17), though the extent and cause of this variation in RpoS between strains is still a matter of some controversy (16, 17).

Microarray studies (18–20) have shown that RpoS controls the expression of at least 500 genes (over 10% of the genome) either directly or indirectly, but the set of RpoS-regulated genes differs across environmental conditions. For most RpoS-controlled genes, it is not known whether the gene is directly transcribed by RpoS or regulated indirectly, as a consequence of RpoS transcribing other genes. Previous studies have not investigated the impact of changes in RpoS levels on the RpoS regulon, or whether quantitative differences in RpoS levels between environmental conditions influence the observed differences in what genes are RpoS-regulated. It is clear that *E. coli* has a complicated regulatory network to fine-tune RpoS levels to different conditions, but we do not yet know the consequences of this regulation.

In this study, we tested the hypothesis that members of the RpoS regulon vary in their response to RpoS levels. Using a combination of ChIP-seq and RNA-seq, we identified RpoS-regulated genes, and we showed that genes vary in their sensitivity to RpoS levels in a manner dependent on sequences outside of the core promoter region. Sensitivity of genes to RpoS levels corresponds to the order in which genes are induced during the transition into stationary phase, and genes with different levels of sensitivity are enriched for specific functional groups. Thus, the level of sensitivity of genes to RpoS controls the physiological response to different stress conditions.

## Materials and Methods

### Culture conditions

Cells were grown in 5mL of LB broth (1% tryptone, 0.5% yeast extract, 1% NaCl) in 150 × 18 mm tubes, positioned vertically and shaking at 225 rpm at 37°C in a water bath, unless otherwise specified in the text. When required, antibiotics were used at the following final concentrations: ampicillin at 100 μg/ml (for plasmids) or 25 μg/ml (for chromosomal integration); chloramphenicol at 20 μg/ml; kanamycin at 50 μg/ml.

### Strains and genetic manipulations

Strains and plasmids used in this study are listed in Table S1. The wild-type genetic background for this all experiments except for ChlP-seq is BW27786, a strain designed to give a graded transcriptional response to increasing arabinose concentration (21). To create a strain of this background lacking *rpoS,* the Δ*rpoS746::kan* allele of JW5437 (22) was moved by P1 transduction into BW27786, creating strain DMS2545.

The arabinose-inducible RpoS strain was created by PCR amplifying the *kan* gene and the P_araB_ promoter of plasmid pAH150 (23) using primers ParaBRpoSRecomb-F and ParaBRpoSRecomb-R (Table S2). This PCR product was then integrated into the *nlpD* gene (i.e. 5’ of *rpoS*) in a MG1655 background using plasmid pKD46 (24), and P1 transduced into BW27786, creating strain DMS2564. This strain thus lacks both the native transcriptional and translational control of RpoS.

For flow cytometry experiments, plasmid pDMS123 (25), which contains a *otsB-gfp* transcriptional fusion, was transformed into DMS2564 by the method of Groth et al. (26).

ChlP-seq experiments used strain RPB104, an unmarked derivative of MG1655 that expresses a C-terminally SPA-tagged derivative of RpoS from its native locus. This strain was constructed by P1 transduction of kan^R^-linked *rpoS-SPA* from a previously described strain (27). The kan^R^ cassette was removed using the pCP20 plasmid that encodes Flp recombinase (24).

### Construction and chromosomal integration of lacZ fusions

The *gadB* and *astC* plasmids were built by standard cloning methods. The *gadC* and *astA* promoter regions with transcription factor binding sites were PCR-amplified with primers gadCpromoter +/-and astApromoter +/-(Table S2), which included *KpnI* and EcoRI restriction sites for cloning. Cloning of core promoter regions was performed by annealing oligonucleotides designed to contain the whole RpoS binding region, as predicted by Fraley *et al.* (28) and Castanie-Cornet and Foster (29). Oligonucleotides were annealed by heating 1 μM of forward and reverse primer for one minute at 100°C with 5 mM MgCl_2_ and 7 mM Tris-HCl and then cooling slowly to room temperature. Inserts and plasmid (pLFX) were digested with EcoRI-HF and KpnI-HF (NEB), ligated with T7 ligase (NEB) and cloned into strain BW23473. Transformants were miniprepped and inserts were verified by Sanger sequencing.

The *gadA* and *hdeA* plasmids were built using Gibson assembly (30) with the NEBuilder HiFi Assembly kit (New England Biolabs). The *gadA* and *hdeA* promoter regions were PCR amplified with primers hdeAHiFi+/-and gadAHiFi+/-. The core promoter was cloned with the single long oligo hdeAcoreHiFi or gadAcoreHiFi, as predicted by Arnqvist et al (31) and De Biase et al (32). PCR products or oligonucleotides were mixed with pLFX digested with KpnI-HF and EcoRI-HF, and assembled according to the manufacturers’ instructions. Mixtures were cloned into strain BW23473, transformants were miniprepped, and inserts were verified by Sanger sequencing.

*lacZ* fusion plasmids were integrated into strain DMS2564 with helper plasmid pPFINT (33). Single-copy integrants were confirmed using the PCR assay of Haldimann and Wanner (23).

### Quantitative Western Blotting

Quantitative western blotting was used to measure RpoS levels. Cells were inoculated from frozen cultures into 5 mL of LB, and grown overnight at 37 °C, shaken at 225 rpm. 5 μL of this overnight culture was diluted into 5 mL of LB with the appropriate concentration of arabinose, and grown for 20 h. 100 μL of overnight culture to be assayed was centrifuged and resuspended in 1X Laemmli sample buffer (Sigma-Aldrich) and boiled for 5 minutes. Samples were diluted 1:10 in 1X Laemlli, and 10 μL was electrophoresed on a 10% polyacrylamide gel (Bio-Rad) in tris-glycine running buffer (25 mM Tris base, 250 mM glycine, 0.5% SDS) at 100 V for 90 min at room temperature. Proteins were transferred to an Immobilon-FL PVDF membrane by electrophoresis at 100 V for 45 min at 4 °C in transfer buffer (48 mM Tris base, 39 mM glycine, 20% methanol, and 0.0375% SDS). Membranes were blocked by overnight incubation in Odyssey blocking buffer (Li-Cor) at 4 °C.

The blocked membrane was probed with affinity-purified monoclonal antibodies to RpoS (clone 1RS1) and RpoD (clone 2G10) (NeoClone) at a final concentration of 0.4 μg/mL in Odyssey blocking buffer plus 0.2% Tween 20 for 1 hour at room temperature. The membrane was washed four times for five minutes each with 15 mL of 1X TBST. A fluorescent secondary antibody (IRDye 800CW goat anti-mouse, Li-Cor) was diluted 1:10,000 in a solution of Odyssey Blocking Buffer plus 0.2% Tween 20 and 0.01% SDS and incubated in the dark for 1 hour at room temperature. The membrane was washed as before, dried for 2 hours between sheets of Whatmann 3MM blotting paper, and imaged on a LiCor Clx fluorescent imager.

Band intensity was estimated using Image Studio 2.1 (LiCor). RpoS levels were divided by RpoD levels to normalize for differences in total protein levels. The ratio of RpoS to RpoD is biologically meaningful because this ratio, rather than RpoS level alone, dictates levels of transcription from RpoS-dependent promoters due to sigma factor competition (34).

### RNA-seq experiments and analysis

Cells were inoculated from frozen cultures into 5 mL of LB, and grown overnight. 5 μL of this overnight culture was diluted into 5 mL of LB and grown for 20 hours. (For the intermediate 26% RpoS condition, a final concentration of 10^−4^% arabinose was also added.) RNA was purified from 200 μL of overnight culture by pelleting and resuspending in 500 μL of Trizol at 65 °C, followed by purification on a column (Direct-Zol, Zymo Research). Samples received two 30-minute DNase treatments using TURBO DNA-free (Ambion) following the manufacturer’s instructions. RNA samples were then purified on a column (RNA Clean & Concentrator, Zymo Research). Samples were stored at −80 °C until used. Three samples were prepped from each culture and pooled to generate sufficient RNA. Two biological replicates were prepared for each strain or condition of interest. rRNA depletion, cDNA synthesis, library preparation, and sequencing were performed by a commercial provider (Otogenetics, Norcross, GA). Paired-end, 100 bp sequences were generated for 7-15 million reads per sample.

Before reads were mapped, the first ten base pairs of each read were trimmed using FASTX-Toolkit 0.0.13. Reads were mapped to the NCBI K12 reference genome (NC_000913.2 Escherichia coli str. K-12 substr. MG1655) using BWA v0.7.5a (35).The number of read pairs mapped to each gene was counted with HTSeq 0.6.1 (36). Differential expression analysis was performed with DESeq v2.13 (37). All p-values were first FDR adjusted using the procedure of Benjamini and Hochberg (38). p-values were then further Bonferroni-adjusted for the three comparisons between pairs of RpoS levels. All differential expression p-values reported in this paper reflect both the FDR and Bonferroni-adjustments.

To determine if a gene differed significantly from the null expectation of linearity, we calculated the probability of the observed read count value at 26% RpoS if the true expression level was given by the linear prediction between the endpoints. Note that to calculate both the expected read count and the probability of the observed value under the null (i.e. the p-value) our model required estimating both the DEseq size factor (for scaling) and the dispersion (a variance factor) for each of the samples. The negative binomial probability model (routinely used to measure count data, (e.g. (37)) was used, with the size factors and dispersion estimated from DESeq (37) to calculate the probability of the observed read count at 26% RpoS.

GO term analysis was performed using the topGO package (39) together with the org.EcK12.eg.db annotation package (40) in R 3.1.0. Enrichment was assessed using the weight01 algorithm (39) together with the Fisher’s-exact test. The GO hierarchy was pruned to include only nodes with at least five associated genes, as significance tests can be unstable for GO terms with fewer genes (39).

A Venn diagram of the number of significant genes in each condition was prepared with EulerAPE 3.0.0 (41).

Analysis of transcription factor binding site enrichment used data from RegulonDB (42). We divided RpoS regulated transcriptional units in RegulonDB into sets: sensitive, insensitive and linear. Then, for each transcription factor in the database we determined how many of the transcription units it regulates fall into each set. We compared this with the number of regulated transcription units within the whole RpoS regulon. A case of enrichment is where a transcription factor regulates a disproportionately large number of units in a particular set (e.g. the sensitive set) compared with what we would expect based on the RpoS regulon as a whole. We identified such cases using a one-tailed hypergeometric test. All p-values were then FDR adjusted using the procedure of Benjamini and Hochberg (38). Transcription factors that with an FDR-adjusted p-value < 0.05 that regulated at least two transcription units in the gene set were considered enriched.

### ChIP-seq analysis

*E. coli* strain RPB104 (MG1655 with C-terminally SPA-tagged *rpoS*) was grown overnight in M9 minimal medium with 0.4% glycerol at 30 °C and then subcultured 1:100 in the same medium, and grown for 60 hours to saturation (OD_600_ of ∼3). ChIP-seq using the M2 monoclonal anti-FLAG antibody was performed as described previously (43). Regions of enrichment (“peaks”) were identified as described previously (44). Relative enrichment was reported as a “Fold Above Threshold” (FAT) score.

We used MEME-ChIP (Version 4.11.2; default parameters) to analyze enriched regions identified by ChIP-seq (45) Regions within 100 bp were merged. The reported sequence motif was identified by MEME (Version 4.11.2) (46), run within the MEME-ChIP environment.

We used MEME (46) within the MEME-ChIP environment (Version 4.11.2; default parameters except that only the given strand was analyzed) to identify enriched sequence motifs in regions surrounding TSSs associated with RpoS ChIP-seq peaks. We analyzed sequences from −45 to +5 relative to each TSS.

We identified directly RpoS-transcribed genes by requiring that (i) the gene start is within 300 bp of a ChIP-seq peak, (ii) the gene is positively regulated by RpoS, as determined by RNA-seq, (iii) no other positively regulated gene starts within 300 bp of the ChIP-seq peak, (iv) there is no associated sequence motif or TSS (identified in (47)) that would be consistent with transcription in the opposite orientation. For regulated genes, we determined whether other genes in the same operon were also regulated, using a published operon list (48) for *E. coli.* Peaks were associated with transcription start sites from a previous study (47) by identifying transcription start sites within 20 bp of a peak.

### QPCR

RNA was isolated as for RNA-seq, except that samples underwent three 30-minute DNase treatments using TURBO DNA-free (Ambion). cDNA was made from the RNA samples using SuperScript VILO master mix (Invitrogen) and stored at −20 °C for use in quantitative real-type PCR (qPCR) reactions.

qPCR was performed using Power SYBR Green master mix (Invitrogen), 2 μL of cDNA, and primers at 300 nM. Cycling was performed at 95 °C for 10 min; and 40 cycles of 95 °C for 15 sec, 58 °C for 15 sec, and 68 °C for 30 sec. Three control genes (*ftsZ, pgm,* and *hemL*) were used in addition genes of interest. These genes were selected using the approach of Vandesompele et al. (49). Details of this selection process, including the other seven genes tested, are available in Supplementary Text 1, Table S3, and Figure S1.

For each gene, a standard curve was made using the following amounts of genomic DNA: 200 ng, 20 ng, 2 ng, 200 pg, 20 pg, and 2 pg. Genomic DNA was extracted from overnight cultures using a Puregene Kit (Qiagen), following the manufacturer’s instructions. Expression levels for each gene were interpolated from the standard curve. Expression levels of experimental genes were divided by the geometric mean of the three control genes.

Statistical assessment of sensitivity was performed by a bootstrapping approach. Bootstrapping, rather than a parametric approach, was appropriate because the data did not conform to parametric assumptions. In this approach, the unit of resampling was the RNA isolated from the 0%, 26% and 100% RpoS conditions on an individual day. For each resampled data set, the median expression at 0% and 100% RpoS was calculated. The linear fit from the 0% RpoS median and the 100% RpoS median was then used to predict the level of expression at the intermediate level of RpoS. Repeating this re-sampling of the data 10,000 times yielded a 95% confidence interval for 26% RpoS. The observed median for 26% was then compared to the confidence interval to test for significance.

### Beta-galactosidase activity

Beta-galactosidase activity was measured using the method of Miller (50). With lacZ fusion strains, the level of sensitivity was quantified, rather than only categorized as sensitive, linear, or insensitive. To quantify the level of sensitivity, for each replicate we calculated the distance between the observed expression at the intermediate RpoS concentration and the expected level based on a linear pattern, standardized by the difference in expression between high and low RpoS conditions. Testing if sensitivities were different from zero used a one-sample t-test, with p-values adjusted for multiple comparisons using Holm’s sequential adjustment method (51). Testing if two sensitivities were different used a two-sample t-test with the same method of adjustment for multiple comparisons.

### Analysis of published RNA-seq data

We analyzed published RNA-seq data for wild-type and Δ*rpoS E. coli* over a time-course of growth into stationary phase (48). Using normalized genome coverage information extracted from wiggle files, we calculated relative abundance for all genes at each of the four stationary phase time-points in the growth curves (the last four time-points for each strain). We arbitrarily selected a threshold coverage value of 500 and excluded any genes scoring below this threshold at all four time-points in wild-type cells. This reduced variability associated with low expression levels. We excluded any genes for which coverage at the final time-point was 0 since this would have prevented normalization. We also excluded any gene for which the first stationary phase time-point had the highest expression value of the four time-points for wild-type cells, since RpoS-dependent expression of these genes is likely to be masked by other factors. We then selected genes whose expression we had found to be induced by RpoS, and separated these genes into insensitive, linear, and sensitive classes. We calculated expression levels for each of these genes relative to expression at the final time-point.

### Accession numbers

RNA-seq data were deposited in GEO with accession number GSE87856. ChIP-seq data were deposited in ArrayExpress with accession number E-MTAB-5339.

## Results

### The RpoS regulon in late stationary phase

To understand the role of the RpoS protein in late stationary phase, we used RNA-seq to compare the transcriptome of wild-type and Δ*rpoS* cells. We observed differential expression of 1044 genes (23% of genes) between these two conditions (p < 0.05). Of the 1044 genes whose expression is influenced by RpoS, 605 are upregulated, and 439 are downregulated (Table S4).

Influencing transcription of 23% of the genome could have many potential phenotypic effects. To better understand the function of these genes, we examined which kinds of gene functions, as described by the gene ontology (GO), are more abundant in the regulon than expected by chance. GO enrichment analysis indicates that the RpoS regulon includes many genes involved in metabolic processes (Table 1); 17 of 18 significantly enriched GO terms are metabolic terms. This metabolic reorganization includes the upregulation of genes encoding glycolytic enzymes and pathways for metabolism of L-arginine to glutamine, or from L-arginine to putrescine and then into succinate. RpoS also drives the downregulation of genes involved in the TCA cycle. These patterns of metabolic regulation are very similar to those identified in late stationary phase in *Salmonella enterica* (52). The only significant GO term not explicitly linked to metabolism (GO:0006970, “response to osmotic stress”) also includes metabolic genes, such as *otsA* and *otsB,* that are involved in trehalose biosynthesis.

**Table 1.** Biological processes enriched in the RpoS regulon.

| GO ID | GO Term | Genes in genome | Observed in regulon | Expected in regulon | p-value |
| --- | --- | --- | --- | --- | --- |
| GO:0009063 | cellular amino acid catabolic process | 69 | 38 | 18.5 | 3.5e-07 |
| GO:0006099 | tricarboxylic acid cycle | 20 | 15 | 5.35 | 8.3e-06 |
| GO:0006096 | glycolytic process | 18 | 14 | 4.81 | 8.3e-06 |
| GO:0006094 | gluconeogenesis | 13 | 9 | 3.48 | 0.0016 |
| GO:0009441 | glycolate metabolic process | 6 | 5 | 1.6 | 0.0063 |
| GO:0006090 | pyruvate metabolic process | 21 | 17 | 5.61 | 0.0180 |
| GO:0006970 | response to osmotic stress | 28 | 13 | 7.49 | 0.0190 |
| GO:0006542 | glutamine biosynthetic process | 10 | 6 | 2.67 | 0.0270 |
| GO:0009060 | aerobic respiration | 65 | 33 | 17.38 | 0.0285 |
| GO:0006006 | glucose metabolic process | 36 | 20 | 9.62 | 0.0317 |
| GO:0016052 | carbohydrate catabolic process | 276 | 94 | 73.79 | 0.0326 |
| GO:0009101 | glycoprotein biosynthetic process | 13 | 7 | 3.48 | 0.0342 |
| GO:0019395 | fatty acid oxidation | 30 | 13 | 8.02 | 0.0354 |
| GO:0000162 | tryptophan biosynthetic process | 11 | 6 | 2.94 | 0.0463 |
| GO:0009246 | enterobacterial common antigen biosynthesis | 11 | 6 | 2.94 | 0.0463 |
| GO:0042398 | cellular modified amino acid biosynthesis | 17 | 7 | 4.55 | 0.0472 |
| GO:0009239 | enterobactin biosynthetic process | 6 | 4 | 1.6 | 0.0472 |
| GO:0000270 | peptidoglycan metabolic process | 51 | 14 | 13.64 | 0.0475 |

Central metabolism is not the only phenotype similarly regulated by RpoS in both S. *enterica* and *E. coli.* Other similarities include transcription of genes involved in antioxidant activities, iron regulation, and Fe-S cluster assembly, the upregulation of proteases, and down regulation of porins. As in S. *enterica,* RpoS in *E. coli* influences the expression of many genes encoding other regulatory proteins, including *csrA, arcA, cra, fur, ihfA, hupA* and *hupB.* These proteins regulate phenotypes including carbon storage (CsrA), central carbon metabolism (ArcA and Cra), iron homeostasis (FUR), and also play a central role in structuring the nucleoid (IHF, HU). Not all of the regulation is identical between *E. coli* and S. *enterica,* however. For example, Lévi-Meyrueis *et al.* (52) noted that RpoS appears to direct switching between many pairs of isozymes, where one isozyme is expressed under conditions when RpoS is abundant, and its partner isozyme is expressed when RpoS levels are low. While some pairs show a similar pattern in *E. coli* (such as *tktA/B* and *acnA/B),* others (such as *fumA/fumC*) do not show this pattern. The reason why these enzymes involved in central carbon metabolism might show this pattern is not clear.

### Genome-wide binding profile of RpoS

While RNA-seq identifies which genes are regulated by RpoS, it cannot distinguish between direct and indirect effects. To determine sites where RpoS binds (and hence likely plays a direct role in transcription), we used ChIP-seq to map the association of RpoS across the *E. coli* chromosome during stationary phase growth in minimal medium. To facilitate ChIP, RpoS was C-terminally SPA-tagged at its native locus. We reasoned that RpoS would only be identified with promoter regions since it is likely released from elongating RNAP complexes. We identified 284 peaks of RpoS ChIP-seq signal covering 260 genomic regions (peaks within 100 bp of each other were merged). 217 of the RpoS-bound regions are intergenic, and 67 are located within genes. We reasoned that annotated genes that are transcribed by RpoS would be positioned close to an RpoS-bound region. Consistent with this, 213 RpoS-bound regions are ≤300 bp upstream of an annotated gene start. These 213 regions include 27 that are intragenic. In 79 cases, we observed an RpoS-bound region ≤300 bp from the starts of two divergently transcribed genes.

We used MEME to search for enriched sequence motifs within the RpoS-bound regions. We detected a highly enriched motif in 107 regions (Figure 1A); this motif closely resembles the known −10 hexamer recognized by RpoS (CTAYACT, where the central YA are not as conserved as the other positions (53)). Moreover, occurrences of the motif are positionally enriched with respect to the ChIP-seq peak center (Figure 1B), indicating that the data have high spatial resolution. Note that the motifs tend to be located just upstream of the peak centers (Figure 1B), as we have observed previously for the *E. coli* flagellar Sigma factor, FliA (44). This presumably reflects the fact that the footprint of initiating RNA polymerase associated with RpoS is not centered on the −10 hexamer.

**Figure 1.**
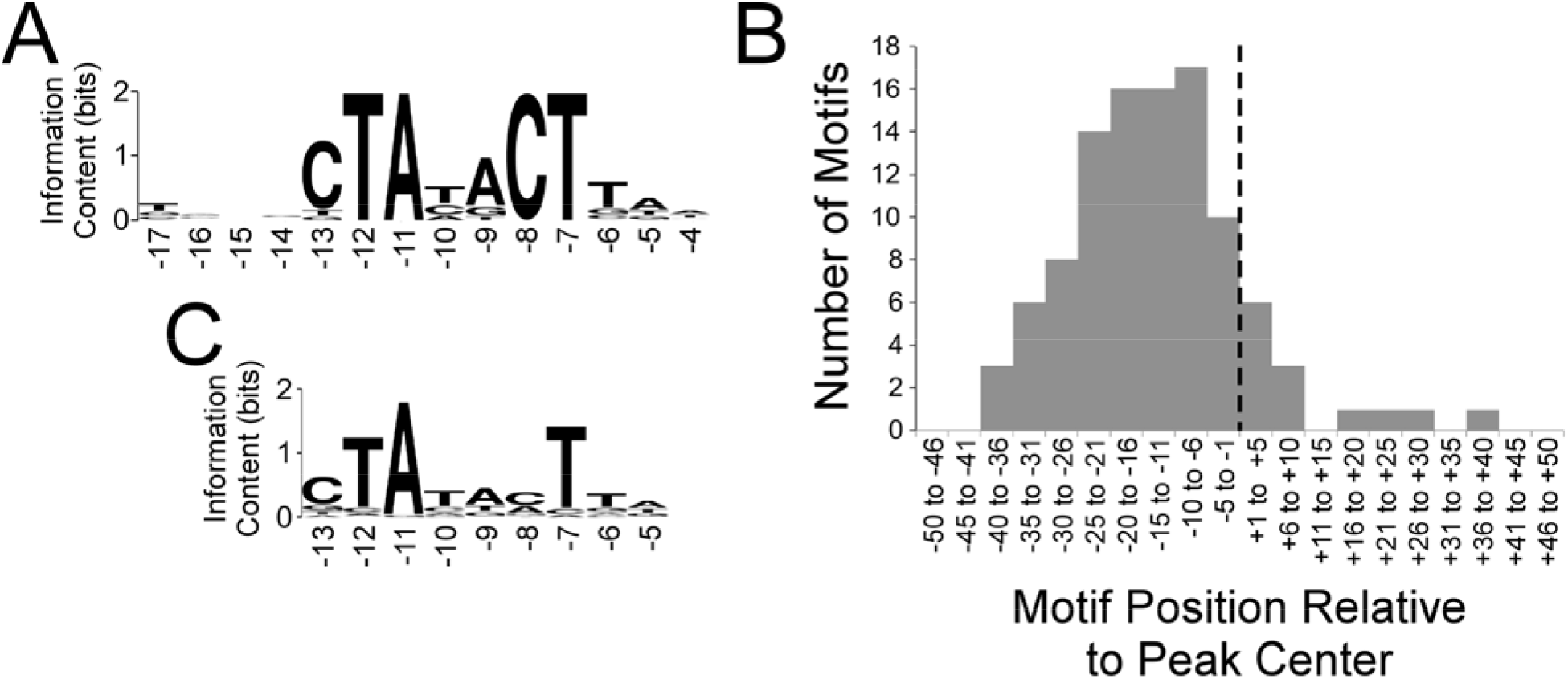
Analysis of RpoS ChIP-seq data. (A) Sequence logo (86) of the enriched sequence motif associated with RpoS-bound regions (MEME E-value 2 × 10^-40^). The sequence logo was generated using WebLogo (87). Numbers below the motif refer to positions relative to the transcription start, based on known sequence preferences for RpoS (53). (B) Histogram showing the frequency distribution of distances between identified motifs and ChIP-seq peak centers. The black, vertical, dashed line indicates the peak center position. (C) Sequence logo (86) of the enriched sequence motif associated with TSSs located within 20 bp of RpoS ChIP-seq peak centers (MEME E-value 3 × 10^-56^). Note that only a subset of the motif identified by MEME is shown, but positions outside the shown region all have information content below 0.2. The sequence logo was generated using WebLogo (87). Numbers below the motif refer to positions relative to the transcription start, based on known sequence preferences for RpoS (53).

The identification of RpoS binding sites allowed us to better understand the role of RpoS in both positive and negative regulation. While similar proportions of the RpoS regulon are positively and negatively regulated by RpoS, it is not clear if this is true at the level of direct regulation. Only 19 genes within 300 bp of a binding site are negatively regulated by RpoS. 19 of the 286 such genes is *fewer* than we would expect by chance if binding sites were randomly distributed around the genome, given than 439 of 4513 genes in the genome are negatively regulated. On the other hand, 111 of 286 genes within 300 bp are positively regulated, a highly significant effect (Fisher’s exact test, p < 10^-16^). Thus, the binding profile of RpoS is consistent with direct positive regulation of many genes, but provides no evidence of direct negative regulation.

### Identification of RpoS-transcribed genes and promoters

We combined the ChIP-seq and RNA-seq data to identify genes that are directly transcribed by RNA polymerase containing RpoS. Thus, we identified 123 RpoS-transcribed genes in 99 transcripts (Table S5), and compared them to other published analyses (18–20, 42, 47, 54–56). In some cases, we identified RpoS-bound regions upstream of genes that were not detected as being RpoS-regulated by RNA-seq. These genes may have promoters that bind transcriptionally inactive RNA polymerase (57, 58). (Inactive polymerase could be bound but physically blocked from transcription elongation by the presence of a bound repressor (59), or could be poised, waiting for a signal to transition to transcription elongation (60, 61).) Alternatively, the disparity between RpoS binding and regulation could be explained by differences in growth conditions between the ChIP-seq and RNA-seq experiments, or by the possibility that the C-terminal SPA tag affects the function of the C-terminus of the protein in the response to transcription activators (57, 61).

The high spatial resolution of sigma factor ChIP-seq can facilitate the identification of specific promoters (44, 62) when combined with nucleotide resolution transcription start site (TSS) maps. Using published TSS data from *E. coli* under stationary phase conditions similar to those used for the ChIP-seq experiment (47), we determined all pairwise distances between RpoS ChIP-seq peaks and stationary phase TSSs. We observed a strong enrichment for peak-TSS distances ≤20 bp (Figure S2). We infer that these 112 TSSs are RpoS-transcribed. Consistent with this, the putative RpoS-transcribed TSSs are associated with −10 hexamers that have features expected of RpoS promoters (Figure 1C). In some cases, RpoS promoters were identified ≤300 bp upstream of genes that were not RpoS-regulated in the RNA-seq data set. We presume that these are RpoS-transcribed genes that otherwise escaped detection because of differences in the growth conditions used for ChIP-seq and RNA-seq.

### The transcriptome at three different RpoS levels

This first view of the RpoS regulon considers RpoS as either present or absent. RpoS levels vary continuously across environmental stresses (7), so we sought to better understand how the level of RpoS in the cell influences transcription of target genes. To do this, we placed the *rpoS* gene under the control of the arabinose-inducible promoter P_araB_. This promoter was integrated just upstream of the native *rpoS* gene, placing transcription under the control of arabinose concentration and removing the 5’ region that regulates translation of the native mRNA (4).

To measure the resulting arabinose-induced expression of RpoS, we employed quantitative western blotting. RpoS levels increased with increasing arabinose concentration, from undetectable RpoS levels, to levels similar to those in wild-type cells (Fig 2a). To confirm that expression was graded and not all-or-none in this system (21), we used flow cytometry to measure expression in individual cells. We transformed the arabinose-inducible RpoS strain with the plasmid pDMS123 (25), which contains the RpoS-dependent *otsBA* promoter fused to *gfp.* As expected, *gfp* expression increased with increasing arabinose concentrations, and at each expression level the population was unimodal (Fig 2b).

**Figure 2.**
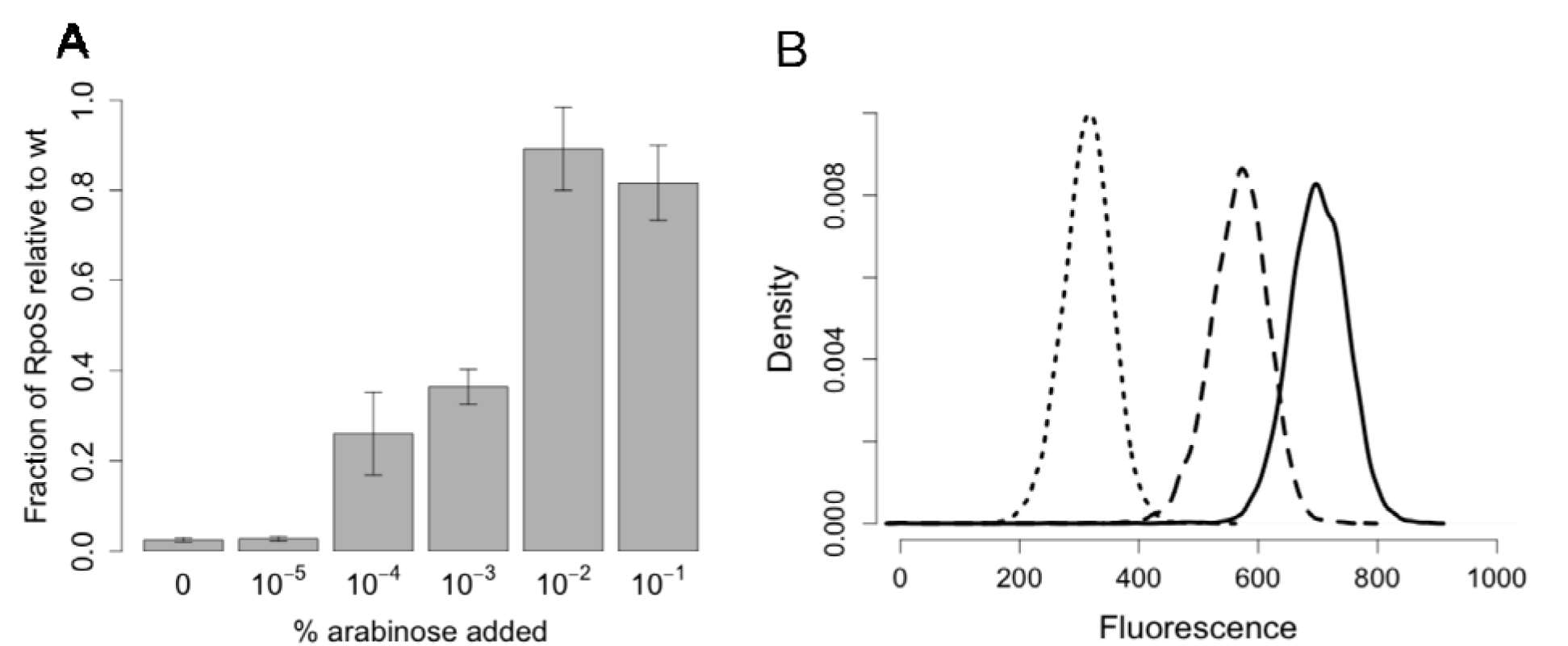
Arabinose control of RpoS protein levels in strain DMS2564. (a) Increasing arabinose results in more RpoS expression in strain DMS2564, as measured by western blotting. n = 4 to 5; error bars represent SEM. (b) Flow cytometric measurement of an *otsB-gfp* fusion under 0%, 10^-4^%, and 10^-1^% arabinose. (Dotted line, dashed line, and solid line, respectively.)

We measured the transcriptome in cells with 26% of wild-type levels of RpoS, achieved with the addition of 10^-4^ % arabinose to cells with our arabinose-inducible *rpoS* strain. Of the genes that are differentially expressed between 26% RpoS and either 0% (ΔrpoS) or 100% (wild-type), 95% are also differentially expressed between 0% and 100% (Fig 3) (p < 0.05).

**Figure 3:**
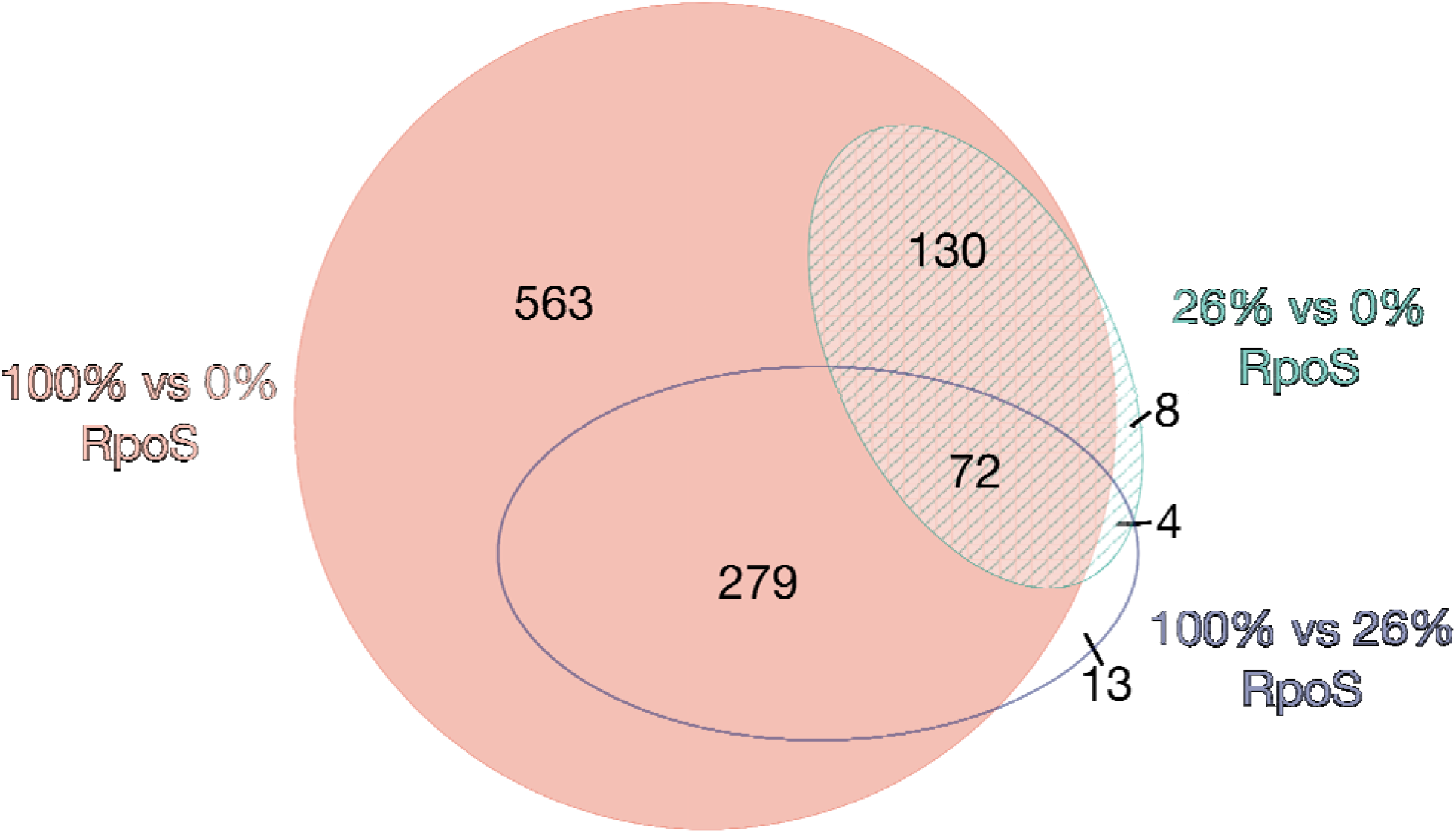
Area proportional Venn diagram showing the number of genes differentially expressed between each of the three conditions.

Nearly all genes that are significantly differentially expressed have monotonically increasing or decreasing patterns of expression between three levels of RpoS. Only two genes (*ytfR* and *ytfT*) have an expression level at 26% RpoS that is significantly higher than the expression at both 100% and 0% RpoS. The only two genes with expression lower in the 26% RpoS condition than either 100% or 0% are *nlpD* and *pcm.* These genes lie immediately upstream of *rpoS. nlpD* was removed from the genome during the construction the arabinose inducible RpoS strain. The *pcm* gene is still present, but the level of transcription was lowered by the genetic modification.

### Classifying quantitative responses to RpoS levels

To explore how genes respond to changing levels of RpoS, we developed a new metric, *sensitivity.* Our null expectation was that gene expression would increase linearly with increasing RpoS concentration. We observed many genes in our RNA-seq data set (such as *osmY*) whose expression at intermediate RpoS levels falls on, or close, to a line drawn between the 0% and 100% RpoS conditions (Fig 4a). We refer to these genes as *linear* in their response to increasing RpoS levels. Other genes (such as *astA*) are transcribed more at 26% than would be expected based on their expression levels at 0% and 100% (Fig 4b). We refer to these genes as *sensitive,* because only a small amount of RpoS results in relatively high levels of transcription. In contrast, some genes (like *gadC*) are expressed at intermediate RpoS levels less than expected based on expression at 0% and 100% RpoS; such genes are referred to as *insensitive* (Fig 4c).

**Figure 4:**
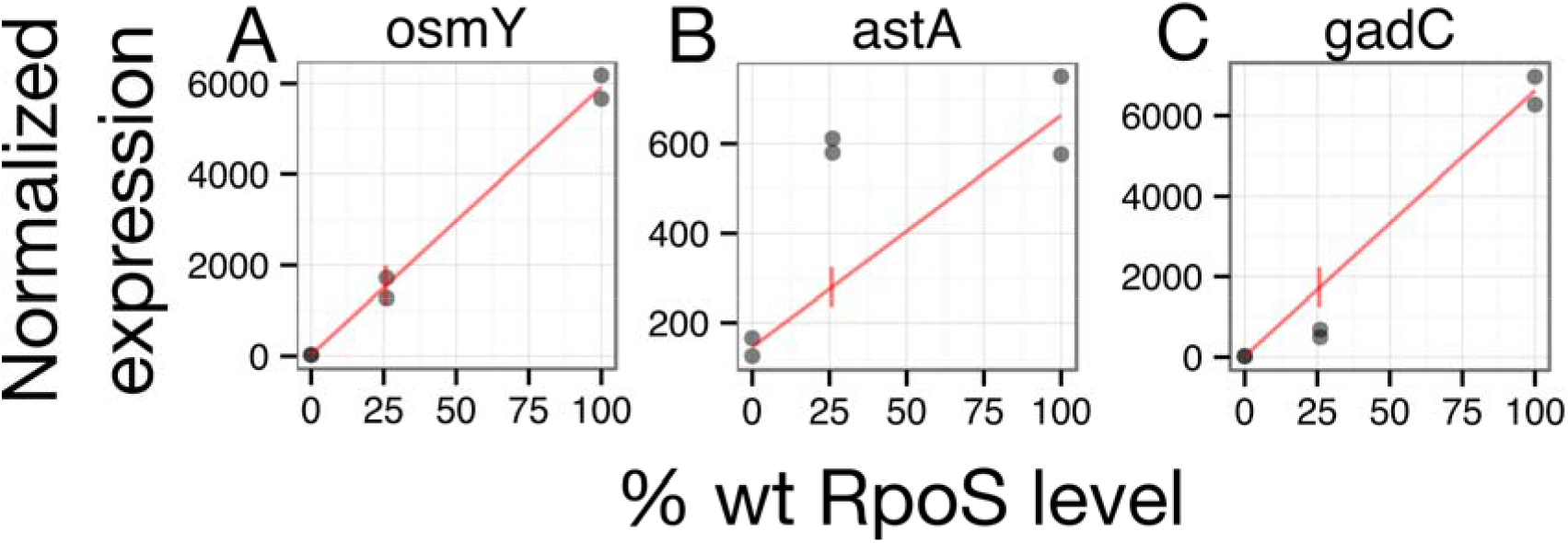
Examples of three classes of patterns of RpoS response. *osmY* expression is linear with RpoS level, *astA* is sensitive to RpoS level, and *gadC* is insensitive to RpoS level. Dots represent normalized expression levels from individual RNA-seq samples. The line is drawn between the mean at 0% and the mean at 100%.

We identified 910 linear, 102 sensitive, and 32 insensitive genes. 96% of sensitive genes and 88% of insensitive genes are positively regulated by RpoS. In contrast, only 53% of linear genes are positively regulated by RpoS, a significant difference (chi-square test, p < 0.001).

To determine whether sensitive or insensitive genes are associated with specific physiological responses to increasing RpoS levels, we again used GO enrichment. We tested the null hypothesis that the functions of these genes are a random sample from the entire RpoS regulon (not the whole genome). The GO terms significantly enriched in the sensitive class are response to osmotic stress, cellular amino acid catabolic process, and fatty acid oxidation (Table 2). Several genes encoding regulators are among the sensitive genes, including *arcA,* which encodes a global regulator of respiratory metabolism, and *rssB,* which encodes the adaptor protein required for degradation of RpoS by ClpXP.

**Table 2.**
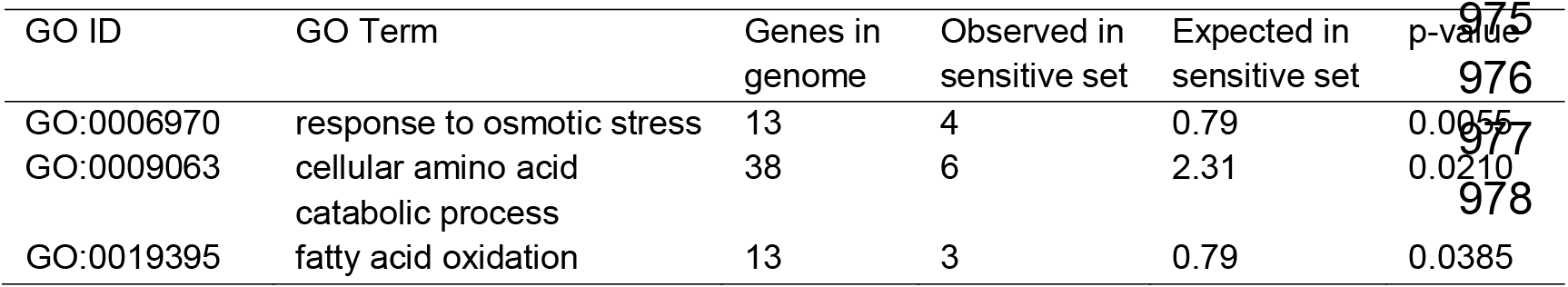
Biological processes enriched in the sensitive genes.

GO enrichment is less useful for understanding the possible function of the insensitive gene set. Three GO terms are enriched (Table 3), but only a few genes with these annotations are present in the insensitive gene set, and their enrichment probably reflects the relatively small number of insensitive genes. More strikingly, the insensitive genes include nearly all of the genes required for acid resistance system 2: the structural genes *gadA, gadB,* and, *gadC,* and the regulator of this system *gadE* (63, 64). In addition, the genes *yhiM, yhiD, hdeA, hdeB, hdeD, mdtE, mdtF,* all of which have been described as having roles in acid resistance (63, 64), are insensitive.

**Table 3.**
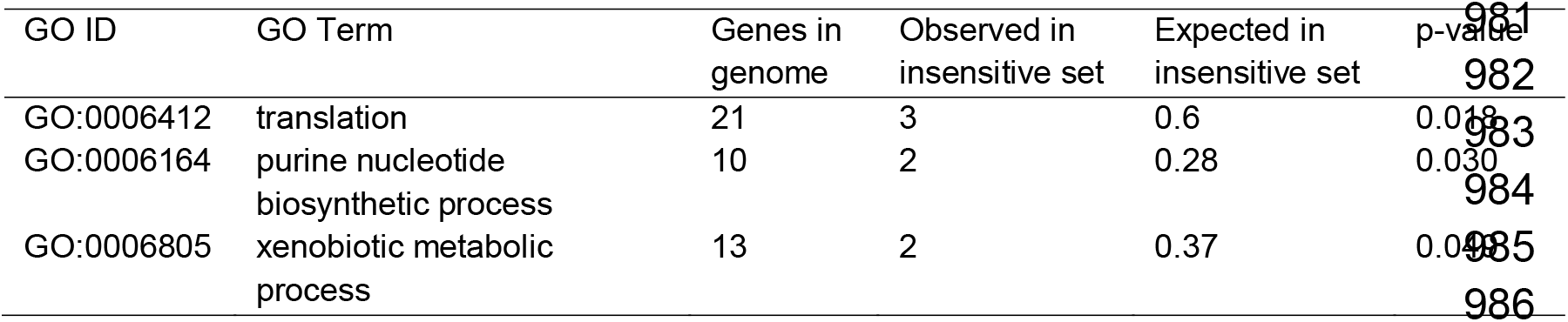
Biological processes enriched in the insensitive genes.

| GO ID | GO Term | Genes in genome | Observed in insensitive set | Expected in insensitive set | p-value |
| --- | --- | --- | --- | --- | --- |
| GO:0006412 | translation | 21 | 3 | 0.6 | 0.018 |
| GO:0006164 | purine nucleotide biosynthetic process | 10 | 2 | 0.28 | 0.030 |
| GO:0006805 | xenobiotic metabolic process | 13 | 2 | 0.37 | 0.044 |

We used reverse transcription coupled to qPCR to confirm the expression patterns of two insensitive genes (*gadC* and *gadE),* and three sensitive genes (*prpR, prpD,* and *astA).* All genes were positively regulated by RpoS (Fig 5), and the median expression at 26% RpoS is consistent with RNA-seq expression patterns for all genes. We used a bootstrapping approach to assess if expression at 26% was significantly above or below the linear expectation at 26% RpoS. The expression of *gadC* was significantly insensitive (p < 10^-4^), and *prpR* expression was significantly sensitive (p < 10^-4^). *astA* expression was marginally significantly sensitive (p = 0.06 for sensitivity), while *gadE* and *prpD* were not significantly different from the linear expectation (p = 0.10 and p = 0.56 respectively).

**Figure 5.**
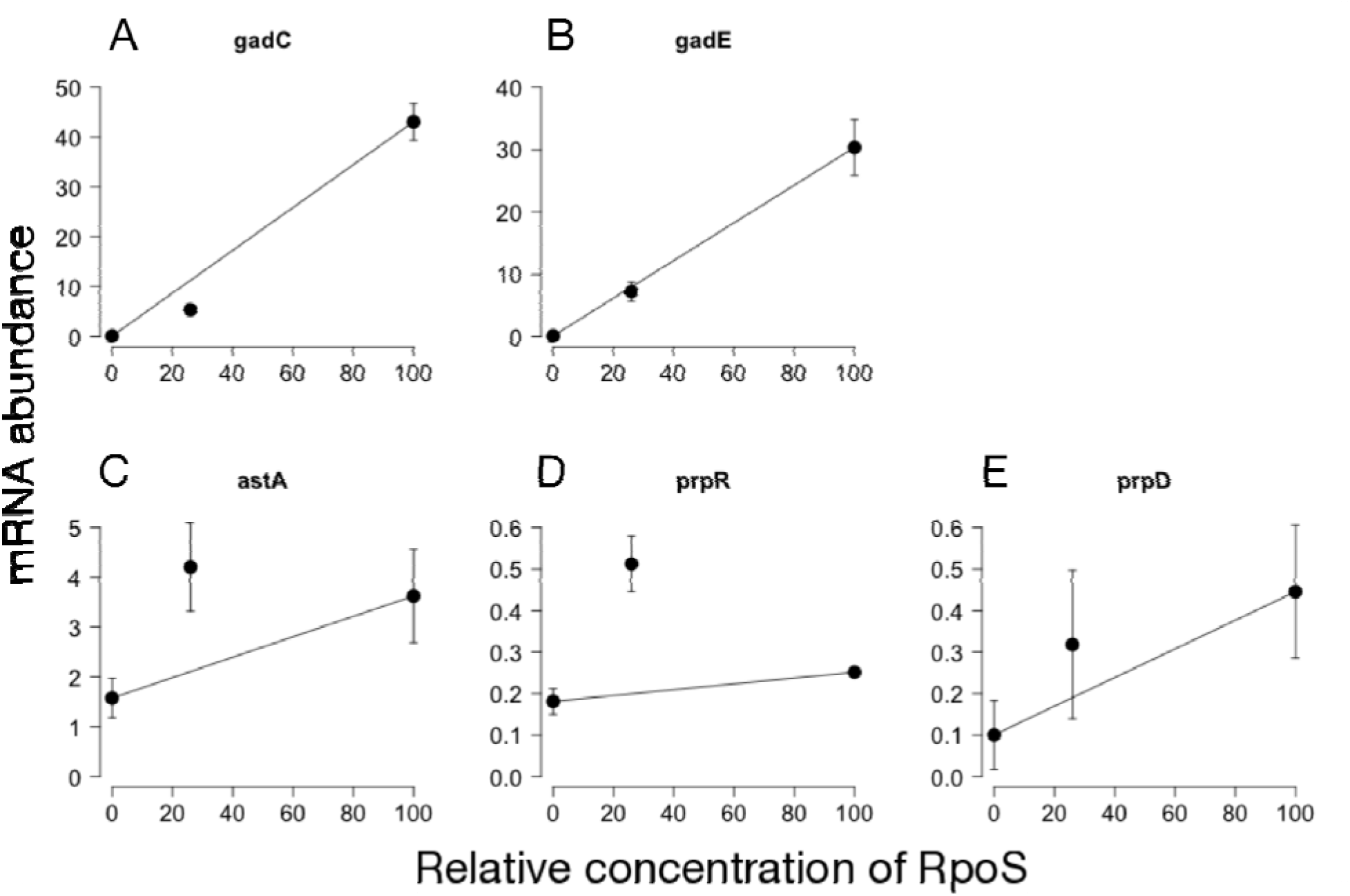
Testing of RNA-seq expression patterns using qPCR. Expression patterns were measured for (a,b) *gadC* and *gadE,* which were insensitive in the RNA-seq data, and (c-e) *astA, prpR,* and *prpD,* which were sensitive in the RNA-seq data. In all cases, the level of transcription at 26% RpoS was on the side of the line predicted by the RNA-seq data, though not always significantly so. *gadC* was significantly insensitive (p < 10^-4^), *gadE* was not significantly different from linear (p = 0.10), *astA* was not quite significantly sensitive (p = 0.06), *prpR* was significant sensitive (p < 10^-4^), and *prpD* was not significant (p = 0.56). n = 6, error bars = SEM.

### Control of sensitivity of expression

What makes one promoter sensitive to RpoS levels and another promoter insensitive to RpoS levels? We hypothesized three possible mechanisms. First, chromosomal location could determine the response to RpoS levels, as is known to occur in the context of total transcription levels (65). Second, it is possible that the DNA sequence of the core promoter drives the response. Finally, it is possible that the binding of transcription factors upstream of the core promoter influences the response to RpoS levels. To test these hypotheses, we cloned the promoters (including all upstream transcription factor binding sites annotated in EcoCyc (66)) of four operons into the *lacZ* fusion plasmid pLFX (33). The four promoters were the sensitive *astCADBE,* and the insensitive *gadA, gadBC,* and *hdeAB-yhiD.* Plasmid pLFX recombines into the lambda-attachment site, placing the fusion in a novel genomic context. While we did not detect binding of RpoS upstream of *astC, gadA, gadB,* or *hdeA* by ChIP-seq, this is likely due to the difference in growth conditions, since these genes have been previously shown to be directly transcribed by RpoS (31, 67, 68).

The pattern of transcription of all four fusions was the same as observed for the respective genes in the RNA-seq data (Fig 6a–d). *astC* transcription was sensitive to RpoS levels (one sample t-test, p = 0.04), while *gadA, gadB,* and *hdeA* transcription were all insensitive (one sample t-test, p = 2 × 10^-6^, p = 10^-5^, p = 0.04 respectively). Since all reporters were placed at the same genomic locus, this result suggests that genomic location is not the determinant of response to RpoS levels.

**Figure 6.**
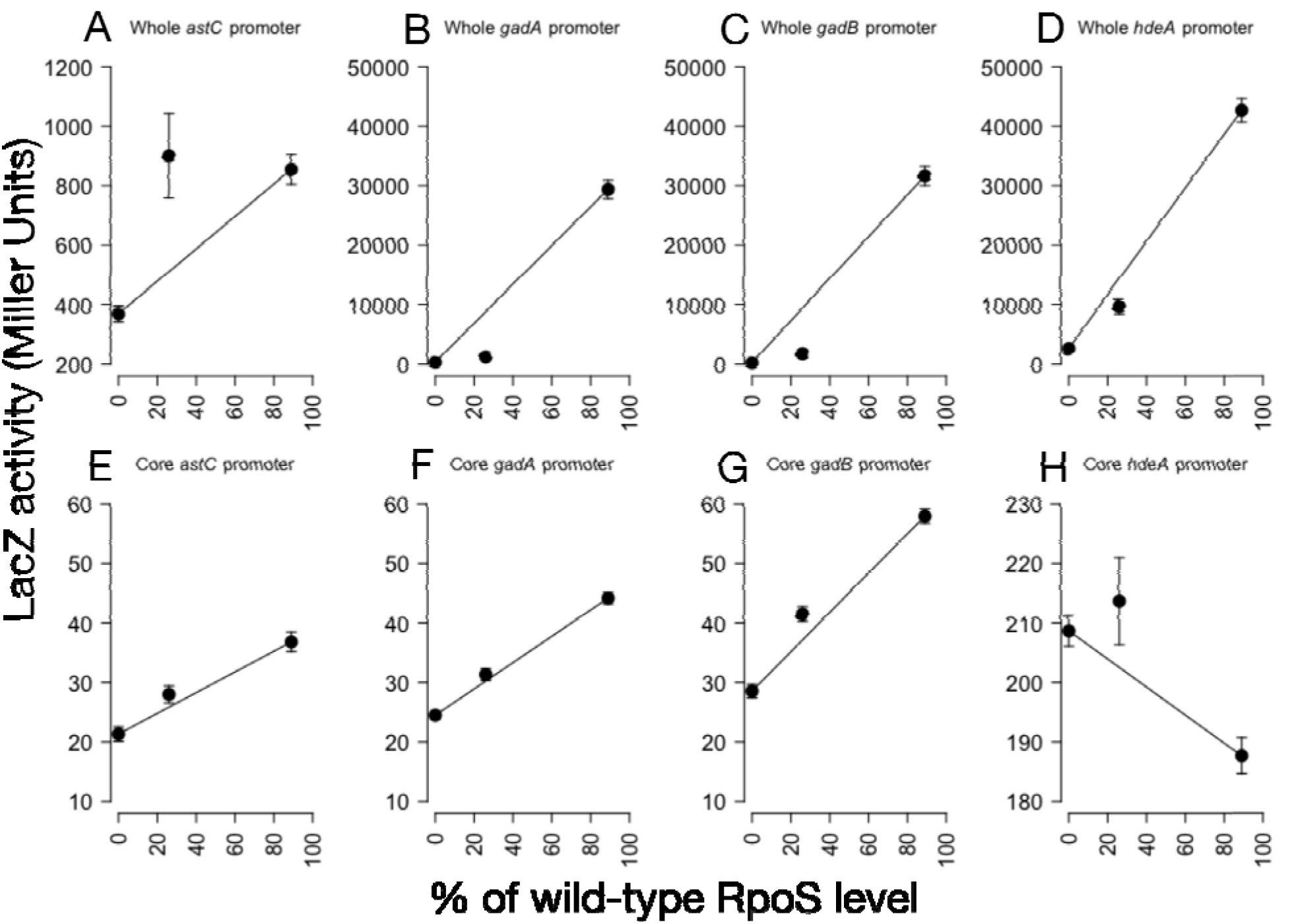
Expression patterns of whole promoters and core promoters only. Expression patterns were measured for the (a-d) whole upstream regulatory region and the (e-h) core promoter only of *astC, gadA, gadB,* and *hdeA.* As expected by RNA-seq data, the whole *astC* fusion was sensitive (p = 0.04, t-test), and the *gadA, gadB,* and *hdeA* fusions were all insensitive (p = 2 × 10^-6^, p = 10^-5^, p = 0.04, all by one sample t-test). The p-values are adjusted for multiple comparisons using Holm’s sequential adjustment method (51). The core promoters (e-h) have much lower levels of maximal transcription than the full length promoters. All four have much altered patterns of sensitivity, although only *gadA* and *gadB* was significantly different from the whole promoter (p = 0.08 (*astC),* p = 0.01 (*gadA),* p = 10^-6^ (*gadB),* p = 0.21 (*hdeA),* all by two-sample t-test with Holm’s correction.) n = 6 to 8, error bars = SEM.

A second potential mechanism to explain the difference between sensitive and insensitive genes is interactions between RpoS and the core promoter sequence. For example, specific nucleotides (or combinations of nucleotides) might tend to confer sensitive or insensitive patterns of transcription. The majority of both sensitive and insensitive genes were not associated with RpoS-bound regions in the ChIP-seq experiment, suggesting that they are indirectly regulated by RpoS (Table 4). The fact that most sensitive and insensitive genes are not bound by RpoS argues against the hypothesis of direct RpoS-DNA interactions driving sensitivity. To see if specific sequence motifs were consistently associated with sensitivity, we used the discriminative motif search feature of DREME (69) to search for motifs that differed between sensitive and linear, or insensitive and linear regulatory sequences. There are 28 ChIP-seq peaks associated with an operon with at least one sensitive gene, and 4 ChIP-seq peaks associated with an operon with at least one insensitive gene. We found no motifs that distinguished these sets of sequences, although the small number of sequences would have restricted the power of such a test.

**Table 4.** Contingency table of the sensitivity of a gene and if is promoter is or is not bound by RpoS.

|  | Sensitive | Linear | Insensitive |
| --- | --- | --- | --- |
| Bound by RpoS | 31 | 87 | 5 |
| Not bound by RpoS | 71 | 823 | 27 |

To directly test the hypothesis that the core promoter region of RpoS-transcribed genes is responsible for determining the sensitivity to RpoS levels, we cloned this short region from the *astC, gadA, gadB, hdeA* promoters into pLFX and recombined the plasmids into the chromosome. These core promoters had absolute levels of transcription much lower than the entire promoter that included upstream transcription factor binding sites (Fig 6e–h). The core promoter sequence for *astC* alone was somewhat less sensitive to RpoS levels than the full-length construct (two sample t-test, p = 0.08), although not significantly so, probably due to the variability of expression from the full-length *astC* reporter. The *gadA* and *gadB* core promoters differed significantly from their full-length promoters (two sample t-test, p= 0.01 and p < 10^-6^, respectively). The *hdeA* core promoter (Fig 6h) is not RpoS-dependent, showing a decline in expression of approximately 10% in the presence of RpoS. This is consistent with the previous finding that the ability of RpoD (but not RpoS) to transcribe *hdeA* is repressed by H-NS protein bound upstream of the promoter (70). The core promoter construct lacks the native H-NS binding sites upstream, and so the selectivity is apparently lost. Thus, the core promoters do not replicate the RpoS sensitivity of their whole promoter sequences. In addition, the three RpoS-dependent core promoters (Fig 6e–g) do not differ from each other in sensitivity (ANOVA, p = 0.32). We conclude that the core promoter is not responsible for RpoS sensitivity.

The *lacZ* fusions suggest that neither genomic location nor core promoter sequence influence the sensitivity of a promoter. The remaining possible mechanism is the binding of specific TFs. If this is the case, we might expect that sensitive and insensitive genes are enriched for binding by different TFs. We looked for such enrichment and found that the sensitive genes are enriched for binding by ArgR, Nac, and NtrC (FDR < 0.05). The insensitive genes are enriched for binding by ArcA, FliZ, GadE, GadW, GadX, H-NS, PhoP, RcsB, and TorR. This very large set of regulators occurs largely because these proteins are all annotated as regulating some or all of the operons involved in AR2: *gadA, gadBC, gadE, hdeAB-yhiD,* and *hdeD.* The action of GadE, GadW, and GadX occurs primarily at these loci, while the other regulators have many additional known binding sites that are not near the promoters of insensitive genes. This specific enrichment for TFs highlights proteins that may be responsible for the sensitive or insensitive patterns of expression.

### Sensitivity to RpoS determines the timing of induction during entry into stationary phase

We hypothesized that the degree of sensitivity to RpoS could impact the timing of expression under conditions when levels of active RpoS increase, such as during entry into stationary phase. A previous study used RNA-seq to monitor the transcriptome over a time-course of growth, including four time-points in stationary phase (48). We analyzed these data to determine whether insensitive, linear, and sensitive genes show differences in the timing of induction. We selected only those RpoS-induced genes whose transcription increased upon entry into stationary phase. (It total, there were 250 such linear genes, 90 sensitive, and 19 insensitive.) We then determined the pattern of expression for each such gene over four time-points beginning at the onset of stationary phase. Although there is considerable variability in the expression patterns of genes, as a group the three classes show clear differences in the timing of induction (Figure 7a). Specifically, sensitive genes are induced most rapidly, followed closely by linear genes. In both cases, expression peaked early in stationary phase and fell between 30 and 180 minutes after entering stationary phase. In contrast, insensitive genes showed relatively little change in expression until the final time-point, 180 minutes into stationary phase. To determine the importance of RpoS on the patterns of gene expression, we repeated the above analysis using data generated from a Δ*rpoS* strain. As expected, the large difference in timing between the groups of genes was greatly diminished (Figure 7b). We conclude that the sensitivity of a gene is associated with the timing of expression during stationary phase in an RpoS-dependent manner.

**Figure 7.**
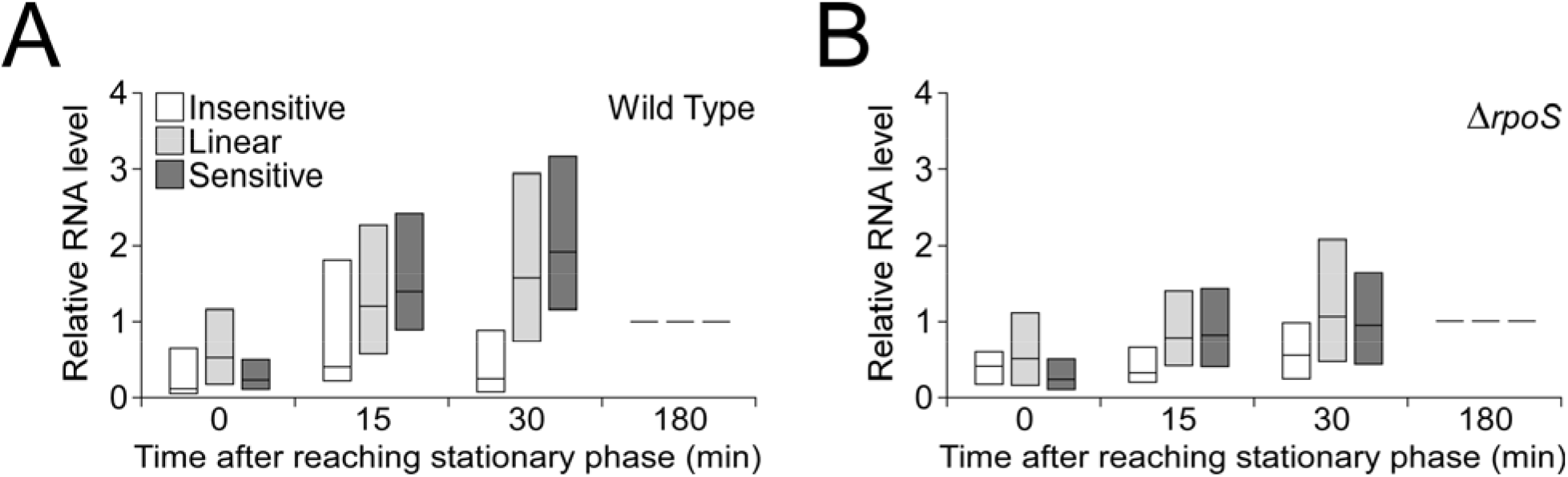
Expression profiles of genes when cells enter stationary phase. **(A)** Relative expression levels of RpoS-regulated genes were taken from a published study (48) for four time-points following entry into stationary phase. All expression values were normalized to those from the final time-point for each gene. The graph shows the range of relative expression values for insensitive (white boxes), linear (light gray boxes), and sensitive (dark gray boxes) genes for each time-point. Boxes represent the 25^th^-75^th^ percentile range, and horizontal lines indicate the median value. **(B)** As above, but for a Δ*rpoS* strain of *E. coli*.

## Discussion

### Expanding the known set of RpoS-transcribed genes and promoters

The *E. coli* RpoS regulon has been widely investigated using targeted and genome-scale approaches. Most genome-scale studies have focused on genes whose expression is altered in the absence of RpoS (18–20, 55, 56). Hence, these studies cannot distinguish between genes that are transcribed by RpoS, and those that are indirectly regulated. ChIP-seq affords a high resolution view of RpoS binding. By combining ChIP-seq with RNA-seq, we have identified 123 RpoS-transcribed genes with high confidence, considerably expanding the known RpoS regulon. Previous studies have suggested a role for RpoS in direct repression of some target genes (47, 57, 71). While we observed negative regulation of 439 genes by RpoS, there were fewer of these repressed genes associated with ChIP-seq peaks than expected by chance, suggesting that direct negative regulation by RpoS is rare.

Only two previous studies have used ChIP methods to map RpoS binding genome-wide in *E. coli.* The first used ChIP-chip to identify 868 RpoS-bound regions (47), many more identified in our study but with considerably lower resolution (median peak length of 324 bp for RpoS ChIP-chip). The second used ChIP-seq, but identified relatively few RpoS-bound regions (54). Of the 63 RpoS-bound regions identified in that study, 41 are shared with those from our study.

The high resolution of ChIP-seq allowed us to identify specific promoter sequences recognized by RpoS. By combining ChIP-seq data with a TSS map, we identified many high-confidence RpoS promoters. These promoters are strongly enriched for the presence of a −10 hexamer, with sequence preferences consistent with several of the previously described features of RpoS promoters (53). Specifically, we observed a preference for a C at position −8 (within the −10 hexamer), a C at position −13 (immediately upstream of the −10 hexamer), and a TAA a positions −6 to −4 (immediately downstream of the −10 hexamer). Previous studies have suggested that RpoS promoters often contain a −35 hexamer (72), although the spacing relative to the −10 hexamer is considerably more variable than for a^70^ promoters. However, we did not detect enrichment of a −35 hexamer-like sequence among RpoS promoters, suggesting that the requirement for this element is weak.

### Sensitivity to RpoS affects timing of expression, and groups functionally related genes

As is true for many transcription factors, RpoS levels vary continuously across a wide range of conditions. Our data show that genes differ in the sensitivity of their response to RpoS levels. Moreover, whether a gene is sensitive or insensitive to RpoS levels is associated with its function, suggesting a physiological rationale for sensitivity. For example, insensitive genes include many of those involved in the glutamate-dependent acid resistance 2 system (AR2). These genes are of particular interest because AR2 allows *E. coli* to survive a pH of 2, an important trait for its ability to pass through the stomach and colonize the gastrointestinal tract (73, 74). To some extent, the shared sensitivity of functionally related genes can be explained by operon structure, i.e. co-transcription of multiple functionally related genes from a single promoter. However, the phenomenon of shared sensitivity for functionally related genes extends beyond operons. For example, the insensitive genes involved in AR2 are transcribed from at least five different promoters (66).

The fact that functionally related genes often have similar patterns of sensitivity to RpoS suggests that sensitivity can serve as a mechanism to control the timing of gene expression, and hence to coordinate specific cellular processes as part of a response to environmental stresses. Consistent with this idea, we have shown that the sensitivity of genes to RpoS levels correlates with the timing of their expression. RpoS sensitivity may drive similar patterns of expression in response to other stresses. Different environmental stresses are known to upregulate RpoS to varying levels (8), suggesting that some insensitive genes may only be expressed under certain stresses. In addition to the effects on gene expression, sensitivity to RpoS may also impact the effects of mutations in *rpoS,* which have been seen to evolve in the lab (75, 76). We expect mutations attenuating RpoS to have the strongest effect on insensitive genes, and the weakest effect on sensitive genes.

### Possible mechanisms of RpoS sensitivity

While the connection between RpoS sensitivity and the timing of gene expression is clear, the molecular basis of sensitivity is less so. Our data indicate that the genomic location of these operons does not determine the expression pattern. Moreover, several lines of evidence suggest that direct interactions between RpoS and the core promoter are also not responsible for determining sensitivity. First, analysis of the ChIP-seq data for the sensitive and insensitive genes finds no motif that distinguishes between them. Second, core promoters from both sensitive and insensitive genes do not replicate the pattern of expression of the full-length promoters. Third, the core promoters of a sensitive operon (*astC*) and two insensitive operons (*gadA* and *gadB*) have indistinguishable patterns of sensitivity, suggesting that what was excluded from those constructs (i.e., binding sites of regulatory proteins) determines the shape of the relationship.

Given our finding that core promoter sequences cannot explain the difference in sensitivity between promoters, we suggest that sensitivity is largely due to the action of specific regulatory proteins bound upstream. If this hypothesis is correct, it could also explain the physiological coherence of these groups. For example, many insensitive genes are involved in the AR2 phenotype and are also regulated by GadX, GadW, and GadE (77–79). If one or more of these three regulators was directly responsible for the insensitive pattern of expression, then this could help to explain the physiological coherence of the insensitive group. The sensitive genes, being a larger group, have no obvious single regulator, although relatively little is known about regulators that function in stationary phase.

It is also possible that the physical properties of promoters play a role in this process, either alone or in concert with transcription factor binding. For example, the supercoiling state of promoters influences levels of transcription (80, 81) and has been implicated in regulating RpoS-dependent transcription (82, 83). It is possible that differential response of promoters to supercoiling levels, either due to interactions directly with RpoS or by changes in the ability of transcription factors to bind (84), plays a role in determining the sensitivity of a promoter. If this type of regulation plays a role, it must be due the structure of the whole promoter itself, rather than supercoiling differences conferred by genomic location (85). We know this because the full promoters cloned into *lacZ* fusions are able to replicate both sensitive and insensitive patterns of transcription, even when moved to the same chromosomal location.

RpoS responds to a wide variety of environmental cues, and regulates genes responsible for many different kinds of responses. This work has demonstrated that one facet of that response, the level of RpoS produced, has varying effects across the entirety of the regulon. The level of RpoS produced in a stress response, together with the action of other transcription factors, may help to tune the RpoS-dependent stress response in ways appropriate for individual stresses.

## Acknowledgements

We thank Rachael Kretsch for experimental help, and Robert Drewell, Xuelin Wu, Jae Hur, Keith Derbyshire and Todd Gray for helpful discussions. This work was supported by HHMI Undergraduate Science Education award #52007544 to Harvey Mudd College and by the NIH Director’s New Innovator Award Program, 1DP2OD007188 (JTW).

## Literature Cited

1. Novick A, Weiner M. 1957. Enzyme induction as an all-or-none phenomenon. Proc Natl Acad Sci U S A 43:553–66.

2. Ptashne M. 2004. A Genetic Switch, Third Edition: Phage Lambda Revisited3rd edition. Cold Spring Harbor Laboratory Press, Cold Spring Harbor, N.Y.

3. Brinsmade SR, Alexander EL, Livny J, Stettner AI, Segrè D, Rhee KY, Sonenshein AL. 2014. Hierarchical expression of genes controlled by the *Bacillus subtilis* global regulatory protein CodY. Proc Natl Acad Sci 111:8227–8232.

4. Battesti A, Majdalani N, Gottesman S. 2011. The RpoS-mediated general stress response in *Escherichia coli.* Annu Rev Microbiol 65:189–213.

5. Hengge R. 2011. Stationary-phase gene regulation in *Escherichia coli,* p. *In* A. Bock, R. Curtiss III, J. B. Kaper, P. D. Karp, F. C. Neidhardt, T. Nyström, J. M. Slauch, C. L. Squires, D. Ussery (eds.), EcoSal-Escherichia coli and Salmonella: cellular and nolecular biology. Washington, DC, ASM Press.

6. Schellhorn HE. 2014. Elucidating the function of the RpoS regulon. Future Microbiol 9:497–507.

7. Lange R, Hengge-Aronis R. 1994. The cellular concentration of the sigma S subunit of RNA polymerase in *Escherichia coli* is controlled at the levels of transcription, translation, and protein stability. Genes Dev 8:1600–1612.

8. Mandel MJ, Silhavy TJ. 2005. Starvation for different nutrients in *Escherichia coli* results in differential modulation of RpoS levels and stability. J Bacteriol 187:434442.

9. Zafar MA, Carabetta VJ, Mandel MJ, Silhavy TJ. 2014. Transcriptional occlusion caused by overlapping promoters. Proc Natl Acad Sci 111:1557–1561.

10. Hengge R. 2009. Proteolysis of sigmaS (RpoS) and the general stress response in *Escherichia coli.* Res Microbiol 160:667–76.

11. Pratt LA, Silhavy TJ. 1998. Crl stimulates RpoS activity during stationary phase. Mol Microbiol 29:1225–1236.

12. Hryckowian AJ, Battesti A, Lemke JJ, Meyer ZC, Welch RA. 2014. IraL is an RssB anti-adaptor that stabilizes RpoS during logarithmic phase growth in *Escherichia coli* and *Shigella*. mBio 5:e01043–14.

13. Chiang SM, Dong T, Edge TA, Schellhorn HE. 2011. Phenotypic diversity caused by differential RpoS activity among environmental *Escherichia coli* isolates. Appl Environ Microbiol 77:7915–23.

14. Bhagwat AA, Tan J, Sharma M, Kothary M, Low S, Tall BD, Bhagwat M. 2006. Functional heterogeneity of RpoS in stress tolerance of enterohemorrhagic *Escherichia coli* strains. Appl Environ Microbiol 72:4978–86.

15. Ferenci T, Galbiati HF, Betteridge T, Phan K, Spira B. 2011. The constancy of global regulation across a species: the concentrations of ppGpp and RpoS are strain-specific in *Escherichia coli.* BMC Microbiol 11:62.

16. Snyder E, Gordon DM, Stoebel DM. 2012. Escherichia coli lacking RpoS are rare in natural populations of non-pathogens. G3 GenesGenomesGenetics 2:1341–1344.

17. Bleibtreu A, Clermont O, Darlu P, Glodt J, Branger C, Picard B, Denamur E. 2014. The *rpoS* gene is predominantly inactivated during laboratory storage and undergoes source-sink evolution in *Escherichia coli* species. J Bacteriol 196:4276–4284.

18. Patten CL, Kirchhof MG, Schertzberg MR, Morton R a, Schellhorn HE. 2004. Microarray analysis of RpoS-mediated gene expression in *Escherichia coli* K-12. Mol Genet Genomics MGG 272:580–91.

19. Weber H, Polen T, Heuveling J, Wendisch VF, Hengge R. 2005. Genome-wide analysis of the general stress response network in *Escherichia coli:* sigmaS-dependent genes, promoters, and sigma factor selectivity. J Bacteriol 187:1591–603.

20. Dong T, Schellhorn HE. 2009. Control of RpoS in global gene expression of *Escherichia coli* in minimal media. Mol Genet Genomics 281:19–33.

21. Khlebnikov A, Datsenko KA, Skaug T, Wanner BL, Keasling JD. 2001. Homogeneous expression of the P_BAD_ promoter in *Escherichia coli* by constitutive expression of the low-affinity high-capacity AraE transporter. Microbiology 147:3241–3247.

22. Baba T, Ara T, Hasegawa M, Takai Y, Okumura Y, Baba M, Datsenko KA, Tomita M, Wanner BL, Mori H. 2006. Construction of *Escherichia coli* K-12 inframe, single-gene knockout mutants: the Keio collection. Mol Syst Biol 2:2006.0008.

23. Haldimann A, Wanner BL. 2001. Conditional-replication, integration, excision, and retrieval plasmid-host systems for gene structure-function studies of bacteria. J Bacteriol 183:6384–93.

24. Datsenko KA, Wanner BL. 2000. One-step inactivation of chromosomal genes in *Escherichia coli* K-12 using PCR products. Proc Natl Acad Sci U S A 97:6640–6645.

25. Stoebel DM, Hokamp K, Last MS, Dorman CJ. 2009. Compensatory evolution of gene regulation in response to stress by *Escherichia coli* lacking RpoS. PLoS Genet 5:e1000671.

26. Groth D, Reszka R, Schenk JA. 1996. Polyethylene glycol-mediated transformation of *Escherichia coli* is increased by room temperature incubation. Anal Biochem 240:302–304.

27. Butland G, Peregrín-Alvarez JM, Li J, Yang W, Yang X, Canadien V, Starostine A, Richards D, Beattie B, Krogan N, Davey M, Parkinson J, Greenblatt J, Emili A. 2005. Interaction network containing conserved and essential protein complexes in *Escherichia coli.* Nature 433:531–537.

28. Fraley CD, Kim JH, McCann MP, Matin A. 1998. The *Escherichia coli* starvation gene *cstC* is involved in amino acid catabolism. J Bacteriol 180:4287–4290.

29. Castanie-Cornet M-P, Foster JW. 2001. Escherichia coli acid resistance: cAMP receptor protein and a 20 bp cis-acting sequence control pH and stationary phase expression of the *gadA* and *gadBC* glutamate decarboxylase genes. Microbiology 147:709–715.

30. Gibson DG, Young L, Chuang R-Y, Venter JC, Hutchison CA, Smith HO. 2009. Enzymatic assembly of DNA molecules up to several hundred kilobases. Nat Methods 6:343–345.

31. Arnqvist A, Olsén A, Normark S. 1994. Sigma S-dependent growth-phase induction of the *csgBA* promoter in *Escherichia coli* can be achieved in vivo by sigma 70 in the absence of the nucleoid-associated protein H-NS. Mol Microbiol 13:1021–1032.

32. De Biase D, Tramonti A, Bossa F, Visca P. 1999. The response to stationary-phase stress conditions in *Escherichia coli:* role and regulation of the glutamic acid decarboxylase system. Mol Microbiol 32:1198–211.

33. Edwards AN, Patterson-Fortin LM, Vakulskas CA, Mercante JW, Potrykus K, Vinella D, Camacho MI, Fields JA, Thompson SA, Georgellis D, Cashel M, Babitzke P, Romeo T. 2011. Circuitry linking the Csr and stringent response global regulatory systems. Mol Microbiol 80:1561–1580.

34. Farewell A, Kvint K, Nyström T. 1998. Negative regulation by RpoS: a case of sigma factor competition. Mol Microbiol 29:1039–51.

35. Li H, Durbin R. 2009. Fast and accurate short read alignment with BurrowsWheeler transform. Bioinformatics 25:1754–1760.

36. Anders S, Pyl PT, Huber W. 2015. HTSeq-a Python framework to work with high-throughput sequencing data. Bioinformatics 31:166–169.

37. Anders S, Huber W. 2010. Differential expression analysis for sequence count data. Genome Biol 11:R106.

38. Benjamini Y, Hochberg Y. 1995. Controlling the false discovery rate: a practical and powerful approach to multiple testing. J R Stat Soc Ser B Methodol 57:289–300.

39. Alexa A, Rahnenfuhrer J. 2010. topGO: Enrichment analysis for Gene Ontology. R package version 2.22.0.

40. Carlson M. org.EcK12.eg.db: Genome wide annotation for *E. coli* strain K12. R package version 3.0.0.

41. Micallef L, Rodgers P. 2014. eulerAPE: drawing area-proportional 3-venn diagrams using ellipses. PLoS ONE 9:e101717.

42. Salgado H, Peralta-Gil M, Gama-Castro S, Santos-Zavaleta A, Muñiz-Rascado L, García-Sotelo JS, Weiss V, Solano-Lira H, Martínez-Flores I, Medina-Rivera A, Salgado-Osorio G, Alquicira-Hernández S, Alquicira-Hernández K, López-Fuentes A, Porrón-Sotelo L, Huerta AM, Bonavides-Martínez C, Balderas-Martínez YI, Pannier L, Olvera M, Labastida A, Jiménez-Jacinto V, Vega-Alvarado L, Del Moral-Chávez V, Hernández-Alvarez A, Morett E, Collado-Vides J. 2013. RegulonDB v8.0: omics data sets, evolutionary conservation, regulatory phrases, cross-validated gold standards and more. Nucleic Acids Res 41:D203–213.

43. Singh SS, Singh N, Bonocora RP, Fitzgerald DM, Wade JT, Grainger DC. 2014. Widespread suppression of intragenic transcription initiation by H-NS. Genes Dev 28:214–219.

44. Fitzgerald DM, Bonocora RP, Wade JT. 2014. Comprehensive mapping of the *Escherichia coli* flagellar regulatory network. PLoS Genet 10:e1004649.

45. Machanick P, Bailey TL. 2011. MEME-ChIP: motif analysis of large DNA datasets. Bioinformatics 27:1696–1697.

46. Bailey TL, Elkan C. 1994. Fitting a mixture model by expectation maximization to discover motifs in biopolymers. Proc Int Conf Intell Syst Mol Biol 2:28–36.

47. Cho B-K, Kim D, Knight EM, Zengler K, Palsson BO. 2014. Genome-scale reconstruction of the sigma factor network in *Escherichia coli:* topology and functional states. BMC Biol 12:4.

48. Conway T, Creecy JP, Maddox SM, Grissom JE, Conkle TL, Shadid TM, Teramoto J, San Miguel P, Shimada T, Ishihama A, Mori H, Wanner BL. 2014. Unprecedented high-resolution view of bacterial operon architecture revealed by RNA sequencing. mBio 5:e01442–1414.

49. Vandesompele J, Preter KD, Pattyn F, Poppe B, Roy NV, Paepe AD, Speleman F. 2002. Accurate normalization of real-time quantitative RT-PCR data by geometric averaging of multiple internal control genes. Genome Biol 3:research0034.

50. Miller JH. 1992. A short course in bacterial genetics. Cold Spring Harbor Laboratory Press.

51. Holm S. 1979. A simple sequentially rejective multiple test procedure. Scand J Stat 6:65–70.

52. Lévi-Meyrueis C, Monteil V, Sismeiro O, Dillies M-A, Monot M, Jagla B, Coppée J-Y, Dupuy B, Norel F. 2014. Expanding the RpoS/σS-Network by RNA Sequencing and Identification of σS-Controlled Small RNAs in *Salmonella.* PloS One 9:e96918.

53. Typas A, Becker G, Hengge R. 2007. The molecular basis of selective promoter activation by the sigmaS subunit of RNA polymerase. Mol Microbiol 63:1296–306.

54. Peano C, Wolf J, Demol J, Rossi E, Petiti L, De Bellis G, Geiselmann J, Egli T, Lacour S, Landini P. 2015. Characterization of the *Escherichia coli σS* core regulon by chromatin immunoprecipitation-sequencing (ChIP-seq) analysis. Sci Rep 5:10469.

55. Lacour S, Landini P. 2004. SigmaS-dependent gene expression at the onset of stationary phase in *Escherichia coli:* function of sigmaS-dependent genes and identification of their promoter sequences. J Bacteriol 186:7186–7195.

56. Dong T, Kirchhof MG, Schellhorn HE. 2008. RpoS regulation of gene expression during exponential growth of *Escherichia coli* K12. Mol Genet Genomics MGG 279:267–277.

57. Rosenthal AZ, Kim Y, Gralla JD. 2008. Poising of *Escherichia coli* RNA polymerase and its release from the sigma38 C-terminal tail for *osmY* transcription. J Mol Biol 376:938–949.

58. Reppas NB, Wade JT, Church GM, Struhl K. 2006. The transition between transcriptional initiation and elongation in *E. coli* is highly variable and often rate limiting. Mol Cell 24:747–757.

59. Schröder O, Wagner R. 2000. The bacterial DNA-binding protein H-NS represses ribosomal RNA transcription by trapping RNA polymerase in the initiation complex1 J Mol Biol 298:737–748.

60. Rosenthal AZ, Hu M, Gralla JD. 2006. Osmolyte-induced transcription:-35 region elements and recognition by sigma38 (rpoS). Mol Microbiol 59:1052–61.

61. Huo Y-X, Rosenthal AZ, Gralla JD. 2008. General stress response signalling: unwrapping transcription complexes by DNA relaxation via the sigma38 C-terminal domain. Mol Microbiol 70:369–378.

62. Bonocora RP, Smith C, Lapierre P, Wade JT. 2015. Genome-scale mapping of *Escherichia coli* sigma54 reveals widespread, conserved intragenic binding. PLOS Genet 11:e1005552.

63. Mates AK, Sayed AK, Foster JW. 2007. Products of the *Escherichia coli* acid fitness island attenuate metabolite stress at extremely low pH and mediate a cell density-dependent acid resistance. J Bacteriol 189:2759–2768.

64. Kanjee U, Houry WA. 2013. Mechanisms of acid resistance in *Escherichia coli.* Annu Rev Microbiol 67:65–81.

65. Bryant JA, Sellars LE, Busby SJW, Lee DJ. 2014. Chromosome position effects on gene expression in *Escherichia coli* K-12. Nucleic Acids Res 42:11383–11392.

66. Keseler IM, Mackie A, Peralta-Gil M, Santos-Zavaleta A, Gama-Castro S, Bonavides-Martínez C, Fulcher C, Huerta AM, Kothari A, Krummenacker M, Latendresse M, Muñiz-Rascado L, Ong Q, Paley S, Schröder I, Shearer AG, Subhraveti P, Travers M, Weerasinghe D, Weiss V, Collado-Vides J, Gunsalus RP, Paulsen I, Karp PD. 2013. EcoCyc: fusing model organism databases with systems biology. Nucleic Acids Res 41:D605–D612.

67. Kiupakis AK, Reitzer L. 2002. ArgR-independent induction and ArgR-dependent superinduction of the *astCADBE* operon in *Escherichia coli.* J Bacteriol 184:2940–2950.

68. Waterman SR, Small PLC. 2003. Transcriptional expression of *Escherichia coli* glutamate-dependent acid resistance genes *gadA* and *gadBC* in an *hns rpoS* mutant. J Bacteriol 185:4644–4647.

69. Bailey TL. 2011. DREME: motif discovery in transcription factor ChIP-seq data. Bioinformatics 27:1653–1659.

70. Shin M, Song M, Rhee JH, Hong Y, Kim Y-J, Seok Y-J, Ha K-S, Jung S-H, Choy HE. 2005. DNA looping-mediated repression by histone-like protein H-NS: specific requirement of Eσ70 as a cofactor for looping. Genes Dev 19:2388–2398.

71. Levi-Meyrueis C, Monteil V, Sismeiro O, Dillies M-A, Kolb A, Monot M, Dupuy B, Duarte SS, Jagla B, Coppée J-Y, Beraud M, Norel F. 2015. Repressor activity of the RpoS/S-dependent RNA polymerase requires DNA binding. Nucleic Acids Res 43:1456–1468.

72. Typas A, Hengge R. 2006. Role of the spacer between the −35 and −10 regions in sigmaS promoter selectivity in *Escherichia coli.* Mol Microbiol 59:1037–1051.

73. Richard HT, Foster JW. 2003. Acid resistance in *Escherichia coli.* Adv Appl Microbiol 52:167–186.

74. Lund P, Tramonti A, Biase DD. 2014. Coping with low pH: molecular strategies in neutralophilic bacteria. FEMS Microbiol Rev 38:1091–1125.

75. Zambrano MM, Siegele DA, Almirón M, Tormo A, Kolter R. 1993. Microbial competition: *Escherichia coli* mutants that take over stationary phase cultures. Science 259:1757–60.

76. Spira B, de Almeida Toledo R, Maharjan RP, Ferenci T. 2011. The uncertain consequences of transferring bacterial strains between laboratories-*rpoS* instability as an example. BMC Microbiol 11:248.

77. Hommais F, Krin E, Coppée J-Y, Lacroix C, Yeramian E, Danchin A, Bertin P. 2004. GadE (YhiE): a novel activator involved in the response to acid environment in *Escherichia coli.* Microbiol Read Engl 150:61–72.

78. Tramonti A, De Canio M, Delany I, Scarlato V, De Biase D. 2006. Mechanisms of transcription activation exerted by GadX and GadW at the *gadA* and *gadBC* gene promoters of the glutamate-based acid resistance system in *Escherichia coli.* J Bacteriol 188:8118–27.

79. Seo SW, Kim D, O’Brien EJ, Szubin R, Palsson BO. 2015. Decoding genome-wide GadEWX-transcriptional regulatory networks reveals multifaceted cellular responses to acid stress in *Escherichia coli.* Nat Commun 6:7970.

80. Travers A, Muskhelishvili G. 2005. DNA supercoiling-a global transcriptional regulator for enterobacterial growth? Nat Rev Microbiol 3:157–169.

81. Dorman CJ. 2006. DNA supercoiling and bacterial gene expression. Sci Prog 89:151–166.

82. Conter A, Menchon C, Gutierrez C. 1997. Role of DNA supercoiling and RpoS sigma factor in the osmotic and growth phase-dependent induction of the Gene osmE of Escherichia coli K12. J Mol Biol 273:75–83.

83. Bordes P, Conter A, Morales V, Bouvier J, Kolb A, Gutierrez C. 2003. DNA supercoiling contributes to disconnect σS accumulation from σS-dependent transcription in Escherichia coli. Mol Microbiol 48:561–571.

84. Cameron ADS, Dorman CJ. 2012. A fundamental regulatory mechanism operating through OmpR and DNA topology controls expression of Salmonella pathogenicity islands SPI-1 and SPI-2. PLoS Genet 8:e1002615.

85. Dorman CJ. 2013. Genome architecture and global gene regulation in bacteria: making progress towards a unified model? Nat Rev Microbiol 11:349–355.

86. Schneider TD, Stephens RM. 1990. Sequence logos: a new way to display consensus sequences. Nucleic Acids Res 18:6097–6100.

87. Crooks GE, Hon G, Chandonia J-M, Brenner SE. 2004. WebLogo: a sequence logo generator. Genome Res 14:1188–1190.

